# Functional Domains of Substance Use and Trauma: A Systematic Review of Neuroimaging Studies

**DOI:** 10.1101/2023.08.11.552870

**Authors:** Cecilia A. Hinojosa, Siara I. Sitar, Joshua C. Zhao, Joshua D. Barbosa, Denise A. Hien, Justine W. Welsh, Negar Fani, Sanne J.H. van Rooij

## Abstract

In a framework for substance use concerning trauma, Hien and colleagues suggested three domains: reward salience, executive function, and negative emotionality. In this PRISMA-guided systematic review, we explored the neural correlates of these domains in individuals who use substances with or without trauma exposure. We included 45 studies utilizing tasks of interest in alcohol, tobacco, and cannabis use groups.

Greater reward, lesser regulation of inhibitory processes, and mixed findings of negative emotionality processes in individuals who use substances versus controls were found. Specifically, greater orbitofrontal cortex, ventral tegmental area, striatum, amygdala, and hippocampal activation was found in response to reward-related tasks, and reduced activation was found in the inferior frontal gyrus and hippocampus in response to inhibition-related tasks. No studies in trauma-exposed individuals met our review criteria.

Future studies examining the role of trauma-related factors are needed and should explore inhibition- and negative-emotionality domains in individuals who use substances to uncover alterations in these domains that place an individual at greater risk for developing SUD.

## 1. Introduction

A substance use disorder (SUD) develops upon continued use of a substance despite experiencing problems due to its consumption (American Psychiatric Association [APA], 2013). SUD is considered a severe public health crisis. About 20.4 million people in the United States were diagnosed with SUD in the past year (National Survey on Drug Use and Health, 2019).

According to the diagnostic criteria, there are a total of 11 symptoms a person may exhibit, including those associated with impairments in inhibitory control, cravings/urges to use the substance, needing more of the substance to achieve the desired effect (tolerance), and withdrawal effects that can only be improved by taking more of the substance (American Psychiatric Association, 2013). SUD can range in severity from mild (exhibiting two or three symptoms) to severe (exhibiting six or more symptoms). SUDs may be associated with adverse physical and mental health outcomes on the individual level and exert large negative outcomes on the societal level. For example, SUD patients experience an increased risk of suicidal outcomes (Poorolajal et al., 2016) and social, academic, and work impairment (Daley, 2013).

Societally, the US suffers billions in annual productivity loss related to lost wages and productivity, crime, and healthcare expenses due to SUD (Schulte & Hser, 2014). Rates of SUD have dramatically increased following the onset of the COVID-19 pandemic (Volkow & Blanco, 2021), especially in underserved communities (Chacon et al., 2021). During the COVID-19 pandemic, African Americans and Hispanic samples started to use or had an increase in substance use at a prevalence ratio of 2.09 than White Americans in response to mental health issues (Baptiste et al., 2020). Thus, it is vital, now more than ever, to better understand the mechanisms that potentially underlie the development of a SUD.

Trauma exposure is a common risk factor for developing or worsening substance use and SUDs (Berenz et al., 2017; Kaysen et al., 2007; Kilpatrick et al., 1997, 2000). Indeed, trauma contributes to the onset of substance misuse (Khantzian, 1985, 1997), and substance misuse can lead to exposure to more trauma (Windle, 1994). Some individuals who experience trauma may develop posttraumatic stress disorder (PTSD). PTSD is a debilitating disorder with a lifetime prevalence rate of 8% in the general population (Kilpatrick et al., 2013) and is highly comorbid with SUD (Pietrzak et al., 2011). Additionally, up to 15% of individuals who have experienced trauma will exhibit some impairing symptoms of PTSD that do not meet the diagnostic threshold (Bergman et al., 2017). While most individuals who experience a traumatic event will not develop PTSD, trauma exposure is still important to examine as it increases the probability of developing substance use and promotes a greater risk of regular use (Degenhardt et al., 2022).

A recent review by Hien et al. (2021) proposed a translational framework designed to advance the creation of interventions for co-occurring PTSD and SUD through understanding the overlapping neurofunctional mechanisms associated with each disorder. The authors provide an overview of studies examining SUD through the context of trauma, focusing on the co-occurrence of PTSD and SUD (PTSD+SUD). Building upon previous work on Research Domain Operating Criteria (RDoC) (Insel et al., 2010) and its counterpart, Alcohol and Addiction Research Domain Criteria (AARDoC) (Litten et al., 2015), the authors propose four domains that contribute to the co-occurrence of PTSD+SUD – reward, executive functioning, negative emotionality, and social cognition - and provide an overview of studies that assess behavioral deficits and biological alterations associated with each domain separately for PTSD, SUD, and co-occurring PTSD+SUD. One limitation of Hien and colleagues’ review was the lack of an in-depth synthesis of the discussed neuroimaging studies exploring each neurofunctional domain. Given the many new developments in the neural mechanisms of SUD and the breadth of neuroimaging studies on this topic, a review of neuroimaging findings related to the domains of the unifying framework proposed by Hien and colleagues is warranted to help explain the neural mechanisms underlying the development of SUD in trauma-exposed samples.

Our review builds upon Hien et al. (2021) by systematically reviewing and synthesizing neuroimaging studies that focus on three domains that may contribute to the overlapping mechanisms affected in SUD and PTSD – reward, executive functioning, and negative emotionality. We chose not to include the social cognition domain, as there is a dearth of neuroimaging studies focused on social cognition in SUD, and more research should be conducted before preliminary conclusions can be drawn. Within this framework, we focus on both samples of individuals who use substances where trauma exposure is not measured and studies that explicitly measure trauma exposure, enabling us to draw conclusions from and identify knowledge gaps for each population. It is important to note that for the studies that do not directly measure trauma exposure, that does not mean the participants in these studies were not trauma-exposed per se, only that their trauma was not directly measured within the confines of the study. Lastly, previous research has found alcohol, cannabis, and tobacco use to be the most used substances by trauma-exposed individuals (Bhalla et al., 2017). As such, we will focus on these three substances within the review. After reviewing the neuroimaging literature, we suggest future research integrating these findings with trauma.

## 2. Methods

### 2.1 Neuroimaging Techniques

Pronounced advances have been made in technologies that explore human brain structure and function in vivo. The main tool used for all studies included in this review is functional magnetic resonance imaging (fMRI). In a research setting, fMRI is used as a proxy to measure differences in brain function across different conditions and psychopathology. The tasks used in the studies reviewed here fall under event-related and block designs. Event-related tasks were developed to improve the temporal resolution; specifically, the presentation of trials is jittered to model the transient responses of separate trial types (Petersen & Dubis, 2012). Block designs are the simplest designs that measure neural activation across a task block to exhibit a single magnitude of activity across certain trials that reflect both the sustained and transient BOLD activity (Petersen & Dubis, 2012). Statistical contrasts are created to determine brain activity during interest versus control trials.

Though not the focus of the current review, some of the studies included functional connectivity (FC) in secondary analyses. FC simultaneously determines brain activation across multiple regions (Friston, 1994). It is thought that if these brain regions activate simultaneously, there is a temporal relationship and a correlation between their activation. Thus, some studies also sought to determine whether a functional relationship existed in brain regions responsible for our domains of interest.

### 2.2 Pathways to SUD Development

#### 2.2.1 Domain 1: Reward

A reward is a stimulus, object, or event that leads to approach behavior, decision-making, positive emotion, or positive reinforcement (Schultz, 2015). Neuroimaging tasks that probe reward-related neurocircuitry in SUD samples fall under the two broad categories of cue-reactivity and reward-processing tasks.

During cue-reactivity tasks, participants view rewarding stimuli such as attractive facial expressions and stimuli related to their drug of choice than images of non-stereotypical attractive faces or drug-neutral cues, respectively. Stimuli can include images or videos of substance paraphernalia or other cues related to substance use, such as odor. Some studies ask participants to rate their craving for the substance on a Likert Scale once a cue is presented.

During reward-processing tasks, participants make decisions resulting in gaining or losing secondary reinforcers, such as money. For example, participants may be asked to guess an outcome when shown a stimulus. If participants guess correctly, they gain money, but if they guess incorrectly, they lose money. Other constructs that can be measured using reward-processing tasks include the neural correlates of reward anticipation.

##### 2.2.1.1 Regions of Interests (ROIs)

Using contrasts that examine reward versus neutral or gain versus loss trials, the tasks described above have been integral in understanding the brain regions important to the reward neurocircuitry, including the anterior cingulate cortex (ACC), orbitofrontal cortex (OFC), ventral tegmental area (VTA), striatum, amygdala, and hippocampus. The ACC is a region important in encoding the net value of rewards to maximize reward and minimize punishment (Hillman & Bilkey, 2012). The OFC is generally responsible for encoding reward identity (Howard & Kahnt, 2021). The VTA is a subcortical region that releases dopamine in response to a reward and is, therefore, integral to the mesolimbic dopaminergic domain (Cai & Tong, 2022). The striatum has a primary role in releasing dopamine and glutamate to other brain regions. The striatum can be separated into subregions, including the globus pallidus, nucleus accumbens, putamen, and caudate. The amygdala is important in reward learning, specifically in creating associations of reward value with neutral stimuli (Baxter & Murray, 2002). Lastly, the hippocampus establishes associations between reward and the context in which reward is received, attributing memories and learning to reward through retaining high-value events associated with reward. Given the importance of these regions in reward, we will focus on these six ROIs for the reward domain.

#### 2.2.2 Domain 2: Inhibition

Researchers have identified executive function, the cognitive processes important in the self-regulation of behavioral achievements, as a key neurofunctional domain to examine the development and maintenance of PTSD+SUD (Hien et al., 2021) given its important role in either disorder (Aupperle et al., 2012; Koob & Volkow, 2016; Kwako et al., 2016). Our review will focus on inhibition (hereafter, the term “inhibition” will be used instead of executive functioning), a key cognitive process critical for substance use (Bickel et al., 2012). Inhibitory control, or the inability to prevent a behavioral or prepotent response (Murphy & Garavan, 2011), is a major deficit in individuals with SUD (Bickel et al., 2012). Two experimental paradigms have been used to measure inhibition within the scanner, the ‘Go/No-Go’ and ‘Stop-signal’ tasks. During the completion of these tasks, participants are generally asked to make a response via a button press to stimuli of one type and withhold their response to stimuli of another.

##### 2.2.2.1 ROIs

Using contrasts that examine correctly withheld responses versus incorrect responses, the tasks described above are optimal in targeting brain regions responsible for inhibition, including the ventromedial prefrontal cortex (vmPFC), dorsal lateral prefrontal cortex (dlPFC), dorsal anterior cingulate cortex (dACC), inferior frontal gyrus (IFG), striatum, and the hippocampus. The vmPFC is responsible for inhibiting a prepotent response during appropriate situations (Aron, 2007; Liddle et al., 2001) and provides goal-directed feedback to motor areas (Munakata et al., 2011). The IFG is important in alerting and shifting attention during salient events (Criaud & Boulinguez, 2013). The dlPFC is essential in inhibitory control (Miyake et al., 2000), sustained attention, and holding task-relevant information in working memory while presenting distractions (Colombo et al., 2015), including stimulus maintenance and retrieval. The striatum is important in stopping when an inhibitory response is warranted (Aron, 2007; Simmonds et al., 2008). Lastly, the hippocampus is key in context-cued response inhibition (Lee & Byeon, 2014). Given the importance of these regions in inhibition, we will focus on these five ROIs for the inhibition domain.

#### 2.2.3 Domain 3: Negative Emotionality

Researchers have identified negative emotionality, disruptions in reactions to negative emotion, as a key neurofunctional domain to examine the development and maintenance of PTSD+SUD (Hien et al., 2021), given the association of negative emotional states with craving and relapse (Koob & Volkow, 2016; Kwako et al., 2016). Paradigms incorporating stimuli that initiate a physiological response have been used to probe participants’ anxiety. Such tasks include showing participants images of pictures within the scanner that vary in valence (negative, neutral, or positive) and arousal (low arousal - high arousal). Images can include pictures of scenes or facial expressions that exhibit emotions, including happy, neutral, and fear. Emotion regulation tasks used include presenting negative images, each preceding instruction to either look at a picture or think about the negative image in a less negative way. Threat-related tasks involve participants learning to anticipate via a cue when they receive a highly annoying but not painful shock.

##### 2.2.3.1 ROIs

The contrast of interest during these tasks includes fearful versus neutral or positive stimuli to explore neurocircuitry responsible for fear responses or threat initiation. The following regions are integral in threat processing, medial PFC (mPFC), rostral and dorsal ACC, and amygdala. The mPFC is a regulatory brain region responsible for behavioral flexibility, encoding motivated actions, promoting fear extinction, and goal-directed action (Jacobs & Moghaddam, 2021). In an anxious state, it is believed that anxiety may suppress executive cortical control, allowing the amygdala to bias attention toward threats and guide responding to that threat (Eysenck et al., 2007; Bishop et al., 2004). The ACC is separated into many subregions. The rostral ACC (rACC) plays a similar role in inhibitory fear-potentiated responses to the mPFC. The dACC is implicated in anxiety-related responding (Seeley et al., 2007). The amygdala is responsible for our fight, flight, or freeze response to potentially threatening environmental stimuli (Phelps, 2004). Lastly, the hippocampus is central in creating associations with contexts in which threatening stimuli are presented (Phelps, 2004). Given the importance of these regions in threat processing, we will focus on these five regions as the ROIs for the negative emotionality domain.

### 2.3 Eligibility

The systematic review was conducted according to the guidelines set by the Preferred Report Items for Systematic Reviews and Meta-analysis (PRISMA). We aimed to provide an overview of neuroimaging studies in SUD that focus on neural mechanisms in three neurofunctional domains to uncover potential neurobiological pathways underlying the development of SUD post-trauma. We conducted two literature searches. The first search focused on studies that included individuals who use substances to investigate neurobiological mechanisms of the three domains irrespective of trauma exposure. The second search focused on a trauma-exposed individuals who use substances to explore potential specific neurobiological characteristics of the three domains related to trauma exposure. Eligibility criteria for the first literature search included: 1) original empirical reports; 2) published in English; 3) used the following neuroimaging techniques; functional magnetic resonance imaging (fMRI) – with or without blood oxygen level-dependent (BOLD) – and positron emission tomography (PET); 4) included a control group in analyses. Exclusion criteria included: 1) meta-analyses/review articles; 2) studies in languages other than English; 3) studies containing only healthy subjects; 4) studies that did not contain a control group; 5) studies that did not make group comparisons; 6) studies that utilized long-term abstinent experimental groups; 7) studies using adolescent or child samples; 8) experimental group was a treatment-seeking sample; 9) intervention or substance was administered during imaging procedures; 10) structural imaging techniques were used; 11) family history of substance use was explored, but not personal substance use in the experimental group; 12) animal studies; and 13) a valid task that measures any of the three domains was not used. Eligibility and exclusion criteria for the second literature search were like the first, with the inclusion of 14) a group of trauma-exposed individuals who use substances as an eligibility criterion.

### 2.4 Search Strategy

For the first literature search, we used PubMed, Web of Science, and Google Scholar and a combination of the following terms and Booleans in the title/abstract of articles: (inhibition OR impulsivity OR impulsiveness); (reward); (anxiety OR anhedonia OR fear OR threat) AND (substance use disorder OR substance abuse OR substance use OR substance misuse OR alcohol abuse OR alcohol use disorder OR alcohol use OR addiction) AND (neuroimaging OR functional magnetic resonance imaging OR fMRI OR functional MRI OR positron emission tomography OR PET) AND (PTSD OR trauma OR posttraumatic stress disorder). For the second literature search, we used the same databases, including the combination of the above terms, with the addition of: AND (PTSD OR trauma OR posttraumatic stress disorder).

### 2.5 Study Selection

Four independent reviewers (CAH, JZ, JB, SVR) screened study titles and abstracts for inclusion with a consensus on selection criteria. Data extraction forms were extracted by three reviewers (JZ, JB, SS) and checked by two reviewers (CAH, SVR). All reviewers resolved any remaining inconsistencies.

The following information was gathered from the articles; sample size for each experimental and control group, the biological sex breakdown of each group, the mean age for each group, study design and methodology, and direction of activation for the brain regions of interest (i.e., brain areas activated related to the three proposed domains).

## 3. Results

### 3.1 Study Characteristics

Refer to **Figure 1** for the PRISMA diagram with a breakdown of the literature reviewed across all domains for the first literature search. In all, 45 articles met the inclusion criteria outlined above (*n*=1195 substance use, *n*=1291 controls), published from 2001-2022. Of the 45 studies, 30 were specific to the reward literature, nine to the inhibition literature, and six to the anxiety sensitivity literature. One study was included for both reward and inhibition domains (Nestor et al., 2011). For the reward domain, of the 31 studies, 14 focused on alcohol (*n*=412 individuals that use alcohol, *n*=398 controls), 12 on tobacco (*n*=224 individuals that use tobacco, *n*=231 controls), and five on cannabis (*n*=142 individuals that use cannabis, *n*=192 controls). For the inhibition domain of the nine studies, five focused on alcohol (*n*=151 individuals that use alcohol, *n*=168 controls), and four on tobacco (*n*=92 individuals that use tobacco, *n*=86 controls) as the substance of interest. Lastly, for the negative emotionality domain, of the six studies, four focused on alcohol (*n*=131 individuals that use alcohol, *n*=177 controls), one on tobacco (*n*=28 individuals that use tobacco, *n*=28 controls), and one on cannabis (*n*=28 individuals that use cannabis, *n*=23 controls) as the substance of interest.

**Figure 1.**
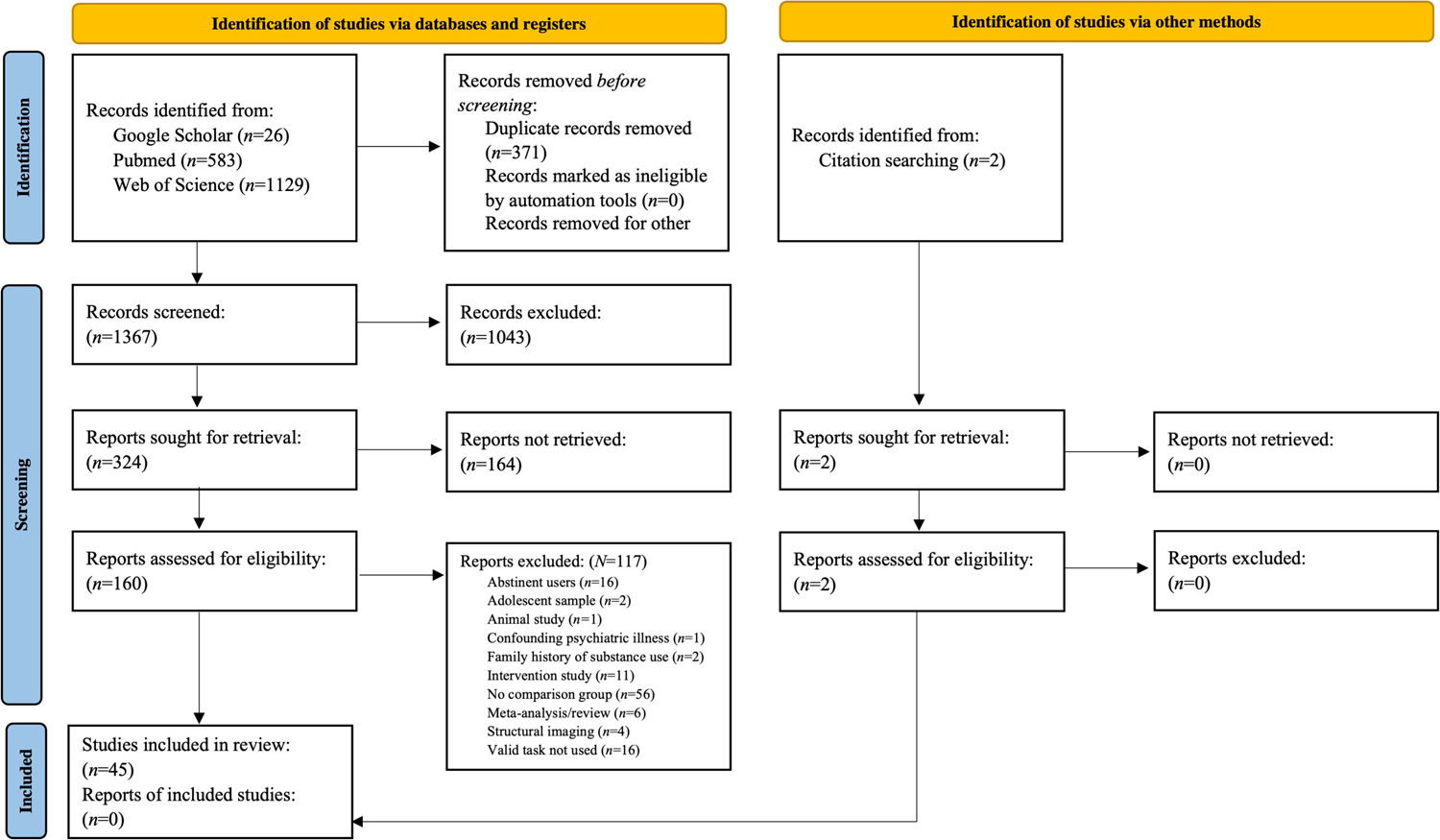
Flow diagram of literature search results from identification to including studies where trauma exposure is not measured.

For studies that contained more than one comparison group (Becker, Gerchen, et al., 2017; Dager et al., 2014; de Ruiter et al., 2009, 2012; Lessov-Schlaggar et al., 2013; Nestor et al., 2011; Nestor et al., 2018; Sjoerds et al., 2014; Strosche et al., 2021; van Hell et al., 2010; Zhou et al., 2019), we report any significant findings between the substance use group and any one of the control groups. One study in the reward domain examined twin pairs discordant for cigarette smoking (Lessov-Schlaggar et al., 2013). For this study, we chose the comparison of individuals who regularly smoked versus those who did not regularly smoke but not their co-twins. One study in the negative emotionality domain included three groups, i.e., individuals with co-occurring alcohol use disorder (AUD)+Anxiety, individuals with AUD-Anxiety, and healthy controls (MacIlvane et al., 2020). To prevent introducing a confound of psychiatric illness in our review, we excluded the findings between the AUD+Anxiety group and the other two groups. Thus, we present only findings from the AUD-Anxiety versus healthy control groups (MacIlvane et al., 2020).

Refer to **Figure 2** for the PRISMA diagram with a breakdown of the literature reviewed across all domains for the second literature search. No articles met the inclusion criteria to be included in the review. A breakdown of reasons we excluded articles is included in **Figure 2**.

**Figure 2.**
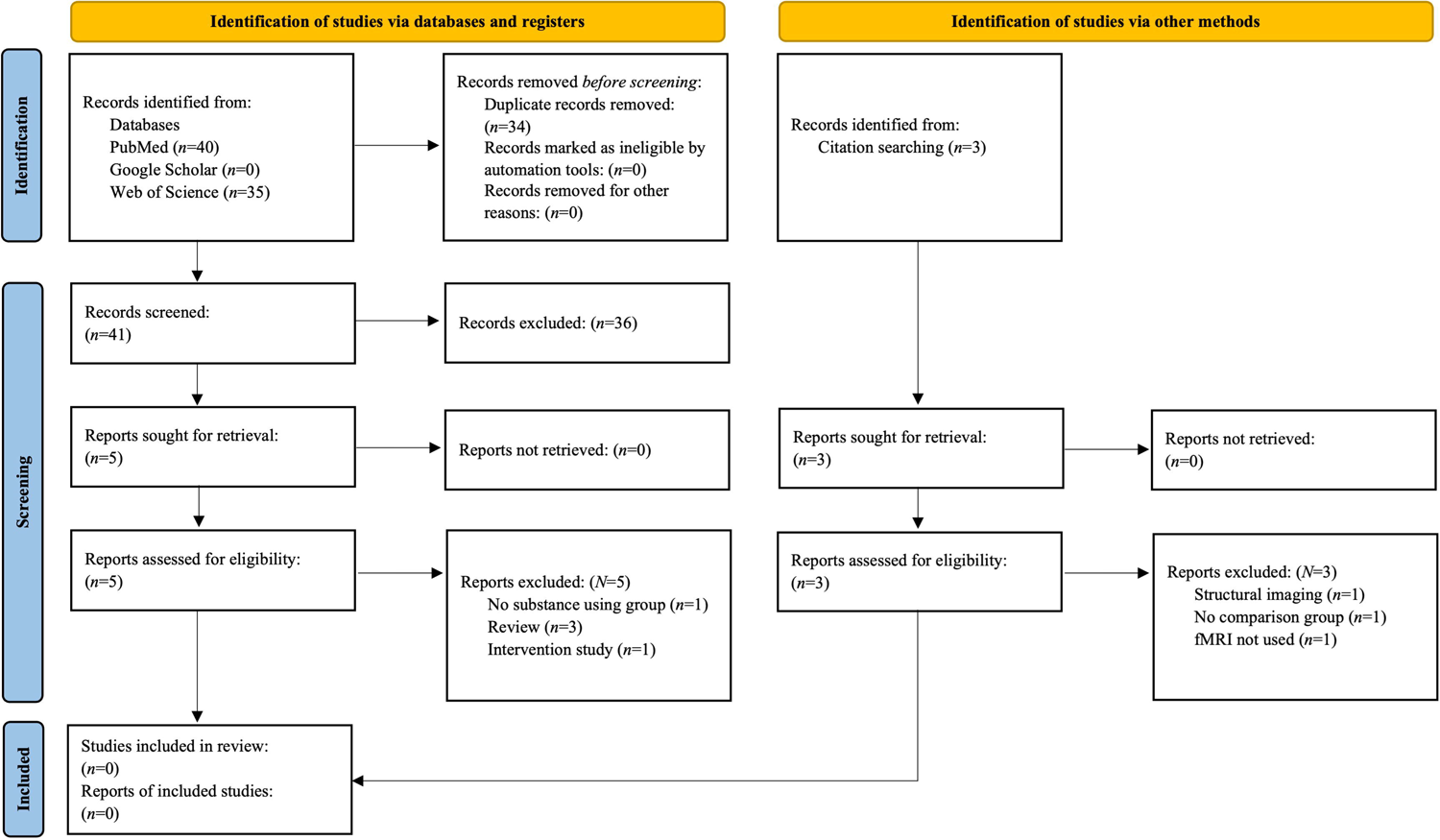
Flow diagram of literature search results for studies where trauma exposure is measured.

### 3.2 Main Findings

A pictorial overview of the main findings can be found in **Figure 3**.

**Figure 3.**
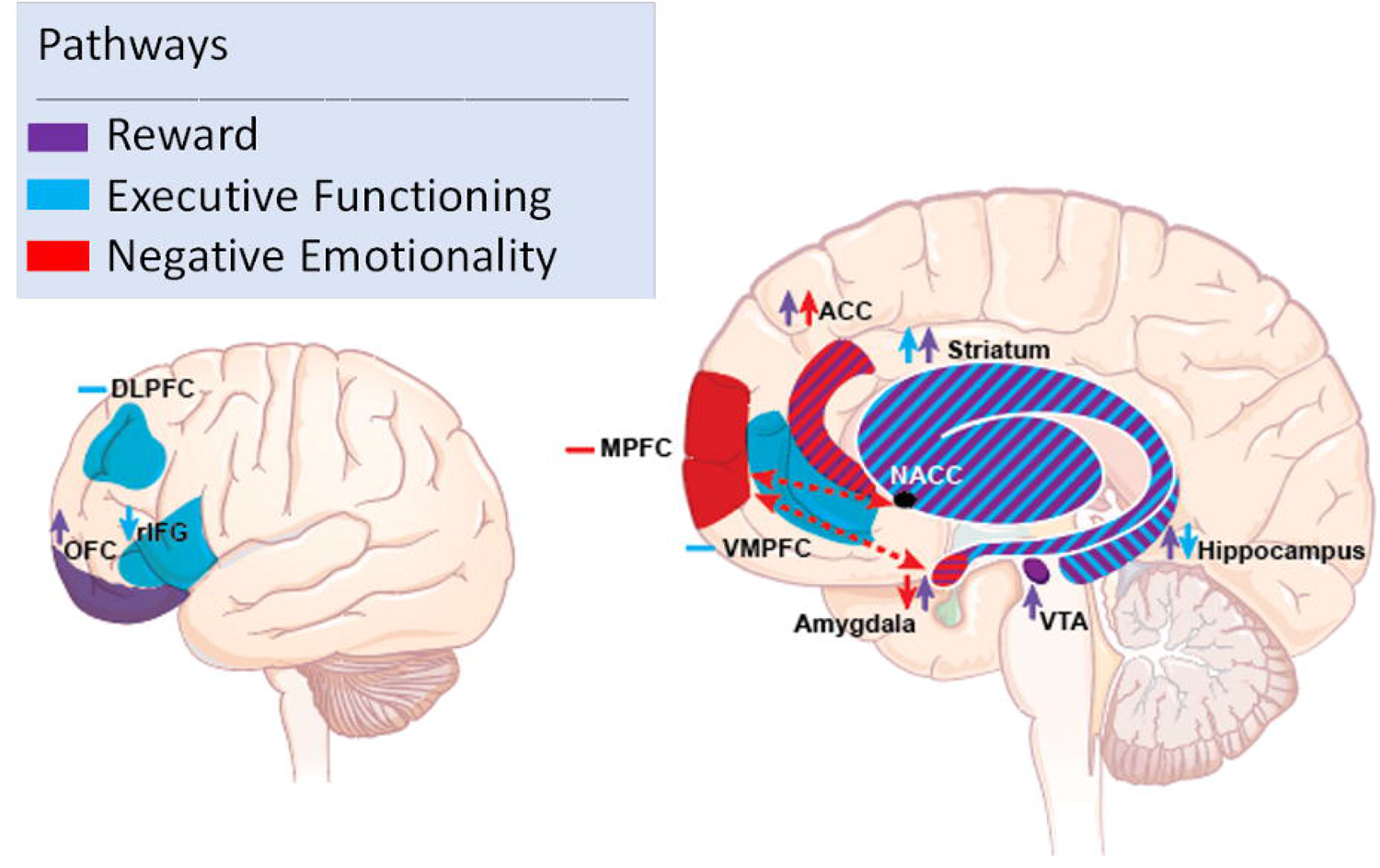
Overview of findings across each domain examined. Purple shading depicts activation found in the reward domain, blue depicts activation found in the executive functioning (inhibition) domain, and red shading depicts activation in the negative emotionality domain. Stripped shading highlights activation found in more than one domain for that respective region. Arrows depict greater or lesser activation in the respective region. Negative signs depict mixed activation found in the respective region. Abbreviations: DLPFC = dorsolateral prefrontal cortex; OFC = orbitofrontal cortex; rIFG = right inferior frontal gyrus; MPFC = medial prefrontal cortex; VMPFC = ventromedial prefrontal cortex; ACC = anterior cingulate cortex; NACC = nucleus accumbens; VTA = ventral tegmental area.

#### 3.2.1 Reward Domain

See **Table 1** for an overview of findings for this domain.

**Table 1.** Reward studies.

| Citation | Substance | Task Used | Experimental Group (n) | Age Mean (SD) | Percentage of Female participants (%) | Control Group | Age Mean (SD) | Percentage of Female participants (%) | OFC | ACC | VTA | Striatum | Hippocampus | Amygdala |
| --- | --- | --- | --- | --- | --- | --- | --- | --- | --- | --- | --- | --- | --- | --- |
| van Holst, Clark, Veltman, van den Brink, Goudriaan, 2014 | Alcohol | Monetary reward task | 19 | 42.5(10.4) | 0% | 19 | 40.4(10.7) | 0% | - | - | - | ↑ | - | - |
| Dager et al., 2014 | Alcohol | Cue-reactivity task | 16 | 18.69 (0.79) | 50% | 13 | 18.54 (0.88) | 61.5% | ↑ | ↑ | - | - | - | - |
| Wiers et al., 2014 | Alcohol | Cue-reactivity task | 20 | 44.30 (7.98) | 0% | 17 | 42.12 (8.3) | 0% | - | - | - | ↑ | - | - |
| Sjoerds et al., 2014 | Alcohol | Cue-reactivity task | 30 | 46.5 (8.5) | 47% | 15 | 46.8 (10) | 27% | ↑ | ↑ | - | ↑ | - | - |
| Becker et al., 2017 | Alcohol | Monetary reward task | 20 | 44.2 (10.3) | 35% | 20 | 44.9 (9.6) | 45% | - | - | - | ↑ | ↑ | - |
| Crane, Gorka, Weafer, Langenecker, de Wit, & Phan, 2017 | Alcohol | Reward guessing game | 27 | 24.00 (2.24) | 44% | 23 | 25.70 (2.95) |  | - | ↓ | - | ↑ | ↓ | ↓ |
| Becker, Kirsch, Gerchen, Kiefer, & Kiefer, 2017 | Alcohol | Monetary reward anticipation task | 31 | 45.4 ± 9 | 22.6% | 35 | 47.7 ± 8.9 | 34% | - | - | - | ↑ | - |  |
| Grodin et al., 2018 | Alcohol | Reward incentive delay with shock task | 21 | 37.33 (13.49) | 38% | 21 | 35.62 (10.96) | 43% | - | ↑ | - | ↑ | - | - |
| Chen et al., 2021 | Alcohol | Pavlovian instrumental transfer task | 94 | 18.4 (0.2) | 0% | 97 | 18.4 (0.2) | 0% | - | - | ↓ | ↓ | - | - |
| Arias et al., 2021 | Alcohol | Number-guessing incentive processing task | 78 | 29.0 (3.5) | 45% | 78 | 28.8(3.7) | 53% | - | ↓ | - | ↑ | - | - |
| Burnette, Grodin, Ghahremani, Galván, Kohno, & Ray, 2021 | Alcohol | Balloon analogue risk task | 16 | 31(9.05) | 31% | 16 | 30.94(10.39) | 31% | - | ↓ | - | - | - | - |
| Strosche, Zhang, Kirsch, Hermann, Ende, Kiefer, & Vollstädt-Klein, 2021 | Alcohol | Cue-reactivity task | 13 | 47 (10) | 39% | 8 | 52 (12) | 50% | ↑↓ | ↓ | ↑ | ↑ | ↑ | ↑ |
| Folco, Fridberg, Arcurio, Finn, Heiman, & James, 2021 | Alcohol | Decision-making task | 15 | 21.20 (2.08) | 100% | 16 | 20.25 (1.57) | 100% | - | - | - | - | - | - |
| Hwang, Martins, Douglas, Choi, Sinha, & Seo, 2022 | Alcohol | Cue-reactivity task | 20 | 31.4 (10.4) | 50% | 20 | 29.4 (9.3) | 50% | - | - | - | - | - | - |
| van Hell et al., 2010 | Cannabis | Monetary reward task | 14 | 24 (4.4) | 1% | 14 | 24 (2.7) | 21% | - | - | - | ↑↓ | ↑ | -- |
| Cousijn et al., 2013 | Cannabis | Monetary decision-making task | 32 | 21.4 (2.3) | 34% | 41 | 22.2 (2.4) | 33% | ↑ | - | - | - | -- | - |
| Filbey et al., 2016 | Cannabis | Cue-reactivity task | 53 | 30.66 (7.48) | 38% | 68 | 31.41 (10.20) | 51% | ↑ | ↑ | ↑ | ↑ | ↑ | ↑ |
| Zhou et al., 2019 | Cannabis | Cue-reactivity task | 18 | 22.94 (2.71) | 0% | 44 | 23.2 (4.32) | 0% | - | - | - | ↑ | - | - |
| Kleinhans et al., 2020 | Cannabis | Cue-reactivity task | 25 | 26.17(4.15) | 48% | 25 | 26.21(5.05) | 48% | - | - | ↑ | ↑ | - | - |
| Yalachkov, Kaiser, & Naumer, 2009 | Tobacco | Cue-reeactivity task | 15 | 27.1 (3.8) | 60% | 15 | 28.7 (6.8) | 60% | - | - | - | ↑ | ↑ | - |
| de Ruiter, Veltman, Goudriaan, Oosterlaan, Sjoerds, & van den Brink, 2009 | Tobacco | Probabilistic reversal-learning task | 19 | 34.8 (9.8) | 0% | 19 | 34.1(9.3) | 0% | - | - | - | - | - | - |
| Zhang et al., 2009 | Tobacco | Backward masking paradigm | 10 | 25.10 (1.07) | 0% | 19 | 23.80 (0.81) | 0% | - | ↑ | - | - | - | ↑ |
| Bühler et al., 2010 | Tobacco | Instrumental motivation task | 21 | 28.0 (4.3) | NR | 21 | 25.7 (6.1) | NR | - | - | - | ↓ | - | - |
| Nestor et al., 2011 | Tobacco | Attentional bias task | 21 | 24.3 (1.2) | 46% | 13 | 23.6 (1.3) | 62% | ↓ | ↓ | - | ↑ | - | - |
| Janes, Nickerson, Frederick, Bde, & Kaufman, 2012 | Tobacco | Cue-reactivity task | 13 | 39.9(10.1) | 100% | 16 | 42.4(10.8) | 100% | - | - | ↑ | ↑ | ↑ | ↑ |
| Lessov-Schlagger et al., 2013 | Tobacco | Adapted card guessing task | 15 | 28.7(3.27) | 100% | 15 | 28.7(3.27) | 100% | - | - | - | ↓ | - | - |
| Nestor, McCabe, Jones, Clancy, & Garavan, 2018 | Tobacco | Monetary incentive delay task | 15 | 23.3 (1.2) | 60% | 15 | 23.8 (1.2) | 47% | ↑ | - | - | ↓ | ↓ | ↓ |
| Duehlmeier & Hester, 2019 | Tobacco | Divergent value learning and errors task | 23 | 25.48 | 35% | 23 | 24.74 | 57% | - | - | - | - | - | - |
| Lawn et al., 2020 | Tobacco | Value-based decision-making task | 19 | 29.5(10.7) | 16% | 19 | 22.7(4.4) | 32% | - | ↓ | - | - | - | - |
| Kunas et al., 2022 | Tobacco | Cue-reactivity task | 38 | 35.18 (10.57) | 55.26% | 42 | 32.36 (10.97) | 73.81% | ↑ | ↑ | - | - | ↑ | - |
| Duehlmeier, Parsons, Malpas, & Hester, 2022 | Tobacco | Feedback learning task | 23 | 25.48 | 35% | 23 | 24.74 | 57% | ↑ | ↑ | - | ↑ | - | - |

*Notes.* Abbreviations: SD = standard deviation; NR = not reported; OFC = orbitofrontal cortex; ACC = anterior cingulate cortex; VTA = ventral tegmental area

##### 3.2.1.1 ACC

Both greater (seven studies) and lesser (six studies) ACC activation in individuals using substances versus controls was observed. This discrepancy is likely dependent on the task used. Specifically, for cue-reactivity tasks, greater ACC was observed in individuals diagnosed with DSM-IV alcohol-dependence (Sjoerds et al., 2014), individuals that drank large amounts of alcohol (Dager et al., 2014), individuals that use tobacco (Kunas et al., 2022), and individuals that use cannabis (Filbey et al., 2016) than their respective control groups during the presentation of their substance cue of interest versus non-substance cue-related images. Conversely, during reward processing tasks, lesser ACC activation was observed in individuals diagnosed with either DSM-IV alcohol abuse or dependence (Arias et al., 2021; Burnette et al., 2021), individuals that experience binge drinking episodes (Crane et al., 2017), and individuals that use tobacco (Lawn et al., 2020; Nestor et al., 2011) than their respective control groups. However, see (Grodin et al., 2018) as this study did not show this pattern. These findings highlight differential mechanisms of the ACC in response to passively viewing reward-related cues versus making conscious decisions regarding reward and loss. Given the role of the ACC in evaluating the value of the reward to maximize value and minimize punishment, individuals that use substances may be unable to correctly categorize rewarding stimuli as rewarding.

Regarding FC, Strosche et al., 2021 found that individuals diagnosed with DSM-IV alcohol dependence exhibited decreased task-related FC between the ACC and insula and ACC and IFG while presenting alcohol-related cues than abstinent individuals (Strosche et al., 2021). During a feedback learning task, widespread greater FC in the dACC and other target regions of interest, including the ACC, was found in individuals who use tobacco (Duehlmeyer et al., 2022). Lastly, one study found that during an unconscious presentation of tobacco smoking images versus the unconscious presentation of neutral images greater FC between the right amygdala and right ACC in individuals who use tobacco than those who did not use tobacco (Zhang et al., 2009).

##### 3.2.1.2 OFC

Greater OFC reward-related activation was observed consistently across studies (eight studies), with a smaller portion showing lesser activation (two studies). In response to cue-reactivity tasks, individuals diagnosed with DSM-IV alcohol dependence (Sjoerds et al., 2014), individuals who transitioned from moderate to heavy drinking (Dager et al., 2014), participants diagnosed with tobacco use disorder (Kunas et al., 2022), and individuals who use cannabis (Filbey et al., 2016) exhibited greater OFC activation than their respective control groups during the presentation of their substance cue of interest versus non-substance cue-related images. During a monetary incentive delay task and feedback learning task, individuals who use tobacco exhibited greater OFC activation than controls (Duehlmeyer et al., 2022; Nestor et al., 2018). During an attentional bias paradigm, individuals who use tobacco exhibited significantly lesser OFC activation than controls (Nestor et al., 2011). In response to an Iowa Gambling Task, individuals who use cannabis showed significantly greater activation in the right OFC during win versus loss trials than controls (Cousijn et al., 2013).

Regarding FC, during an alcohol cue reactivity task, greater FC between the OFC and insula and lesser FC between the OFC and striatum was present in individuals diagnosed with DSM-IV alcohol dependence versus individuals abstaining from alcohol (Strosche et al., 2021). When responding to feedback learning tasks, individuals who use tobacco showed increased connectivity between the ACC and OFC versus those who did not use tobacco (Duehlmeyer et al., 2022).

##### 3.2.1.3 VTA

In response to cue reactivity tasks, VTA activation is consistently greater in individuals with substance use than in controls (four studies), though see (Chen et al., 2021).

Individuals diagnosed with DSM-IV alcohol dependence (Strosche et al., 2021), individuals who use tobacco (Janes et al., 2012), and individuals who use cannabis (Filbey et al., 2016; Kleinhans et al., 2020) exhibited greater activation in the VTA during the presentation of substance-related cues, such as images and odor, than neutral cues. In contrast, individuals at high risk for developing an alcohol use disorder responding to alcohol cues during a Pavlovian instrumental transfer task exhibited decreased activation in the lateral prefrontal cortices than their low-risk counterparts (Chen et al., 2021).

##### 3.2.1.4 Striatum

Overall, individuals who use substances exhibited greater striatal activation than controls (17 studies), though there was variability (five studies showing lesser activation). During cue-reactivity tasks, individuals diagnosed with DSM-IV alcohol dependence (Sjoerds et al., 2014; Strosche et al., 2021; Wiers et al., 2014), individuals with nicotine dependence (Janes et al., 2012; Yalachkov et al., 2009), and individuals who use cannabis (Filbey et al., 2016; Kleinhans et al., 2020; Zhou et al., 2019), exhibited greater striatal activation to their respective preferred substance or other rewarding stimuli than controls. Similarly, using monetary reward and anticipatory tasks (Becker, Gerchen, et al., 2017; Becker, Kirsch, et al., 2017), card guessing tasks (Crane et al., 2017; van Holst et al., 2014), incentive processing tasks (Arias et al., 2021; van Hell et al., 2010), attentional bias paradigms (Nestor et al., 2011), and feedback learning tasks (Duehlmeyer et al., 2022) increased striatal activation is present than controls. During an incentive processing task, individuals with an AUD diagnosis exhibited lower effective directional connectivity between the ACC to the ventral and dorsal striatum than controls without AUD (Arias et al., 2021). A handful of studies have also found decreases in striatal activation during a Pavlovian instrumental transfer (Chen et al., 2021), monetary reward (Nestor et al., 2018; van Hell et al., 2010), card guessing (Lessov-Schlaggar et al., 2013), and instrumental motivation tasks (Bühler et al., 2010) in individuals using substances versus controls.

##### 3.2.1.5 Amygdala

Greater (four studies) and lesser (two studies) amygdala activation was found in response to rewarding stimuli. In cue-reactivity tasks, individuals diagnosed with DSM-IV alcohol dependence (Strosche et al., 2021), individuals who use tobacco (Janes et al., 2012; Zhang et al., 2009), and individuals who use cannabis (Filbey et al., 2016) exhibited greater amygdala activation to their respective preferred substance or other rewarding stimuli than controls. However, using a reward guessing game (Crane et al., 2017) and monetary incentive delay task (Nestor et al., 2018), individuals who use substances exhibit significantly decreased amygdala activation than individuals that do not use substances.

##### 3.2.1.6 Hippocampus

In response to monetary reward tasks and cue reactivity tasks, greater hippocampal activation was found (seven studies), though lesser activation was also found (two studies). Specifically, individuals who use alcohol (Becker, Kirsch, et al., 2017; Strosche et al., 2021), individuals diagnosed with nicotine dependence (Janes et al., 2012; Kunas et al., 2022; Yalachkov et al., 2009), and individuals who use cannabis (Filbey et al., 2016; van Hell et al., 2010) exhibited significantly greater hippocampus activation than individuals who do not use. During a reward guessing game (Crane et al., 2017) and a monetary incentive delay task, (Nestor et al., 2018) decreased hippocampal activation was found in individuals who used substances compared to individuals who did not use them.

##### 3.2.1.7 Summary

Mixed findings appeared in the ACC, whereby greater ACC activation was present in individuals who use substances when passively viewing substance-related cues. However, lesser ACC activation tended to be found during the completion of reward processing tasks, with participants making conscious decisions. This finding could be explained by the inability of individuals who use substances to acknowledge rewarding stimuli as rewarding. Individuals who used substances tended to show significantly greater OFC, VTA, amygdala, and hippocampal activation than controls in response to reward tasks. FC analyses highlight alterations in the ACC and OFC, highlighting FC alterations in more cortical regions than subcortical.

#### 3.2.2 Inhibition Domain

See **Table 2** for an overview of findings for this domain.

**Table 2.** Executive Functioning (Inhibition) studies.

*Notes.* Abbreviations: SD = standard deviation; vmPFC = ventromedial prefrontal cortex; IFG = inferior frontal gyrus; dlPFC = dorsolateral prefrontal cortex
| Citation | Substance | Task Used | Experimental Group n | Age Mean (SD) | Percentage of Female participants (%) | Control Group n | Age Mean (SD) | Percentage of Female participants (%) | vmPFC | IFG | Striatum | dlPFC | Hippocampus |
| --- | --- | --- | --- | --- | --- | --- | --- | --- | --- | --- | --- | --- | --- |
| Ahmadi et al., 2013 | Alcohol | Go/No-Go task | 35 | 18.97 (0.45) | 65.8% | 56 | 18.80 (0.97) | 44.7% | ↓ | - | ↓ | - | ↓ |
| Ames et al., 2014 | Alcohol | Go/No-Go task | 21 | 20.19 (1.4) | 52% | 20 | 20.75 (1.07) | 65% | ↑ | - | - | ↑ | - |
| Campanella et al., 2017 | Alcohol | Go/No-Go task | 19 | 24.7 (3.0) | 58% | 17 | 25.8 (4.2) | 30% | - | - | ↑ | - | - |
| Hu et al., 2016 | Alcohol | Stop signal task | 57 | 29.1 (9.6) | 39% | 57 | 31.5 (11.8) | 46% | ↓ | - | - | - | - |
| Alderson et al., 2021 | Alcohol | Go/No-Go task | 19 | 23.5 (3.1) | 57% | 18 | 25.6 (4.1) | 50% | ↑ | - | - | - | - |
| Nestor et al.,<br>2011 | Tobacco | Go/No-Go<br>task | 10 | 23.0<br>(1.0) | 50% | 13 | 23.6 (1.3) | 62% | ↓ | ↓ | - | ↓ | ↓ |
| de Ruiter et al.,<br>2012 | Tobacco | Stop signal<br>task | 10 | 33.8<br>(9.1) | 50% | 17 | 34.7 (9.7) | 0% | - | ↓ | - | - | - |
| Clewett et al.,<br>2014 | Tobacco | Delay<br>discounting<br>task | 39 | 35.9<br>(8.7) | 33.3% | 33 | 30.1 (7.2) | 47% | - | - | - | - | - |
| Akkermans et<br>al., 2018 | Tobacco | Go/No-Go<br>task | 39 | 22.56<br>(2.84) | 28% | 23 | 21.74<br>(1.82) | 39% | - | - | ↑ | - | - |

##### 3.2.2.1 ACC

In response to inhibition tasks, greater (two studies) and lesser (three studies) activation has been found in individuals who use substances than controls. When completing the Go/No-Go task, individuals who drank heavily (Ahmadi et al., 2013), individuals who drank socially (Hu et al., 2016), and individuals who previously used tobacco (Nestor et al., 2011) exhibited significantly lower ACC activation than controls. Though, the opposite activation pattern has been found using the same task. Specifically, individuals who drank heavily exhibited stronger activation in the right dlPFC and dACC than individuals who drank light during No-Go trials (Ames et al., 2014). These discrepant findings are likely caused by differences in contrasts examined and, therefore, differences in underlying neural processes. Specifically, studies that found lower ACC activation examined the correct inhibition responses versus errors. In contrast, studies that showed greater activation were found using the opposite contrast, error inhibition responses versus correct.

Individuals who experienced binge drinking episodes exhibited greater activation than individuals who drank light in the rACC and increased FC between the rACC and right lateral frontal cortex during the error No-Go > Correct No-Go responses (Alderson Myers et al., 2021).

##### 3.2.2.2 IFG

Two studies showed reduced rIFG activation in individuals who use tobacco. Specifically, significantly lesser activation was found in the IFG during the Go/No-Go task (Nestor et al., 2011) and the Stop Signal Task (de Ruiter et al., 2012) than in controls.

##### 3.2.2.3 dlPFC

Findings of inhibition-related dlPFC activation in substance use were mixed, with one study finding greater and one finding lesser activation. Significantly greater dlPFC activation during the Go/No-Go task was observed in individuals who drank heavily versus individuals who drank light (Ames et al., 2014). Still, less dlPFC activation in individuals who use tobacco than controls, and individuals who used to smoke tobacco were observed using the same task (Nestor et al., 2011).

##### 3.2.2.4 Striatum

Studies showed greater (two studies) and lesser (one study) activation in the striatum during inhibition-related tasks in individuals who use substances. During failed inhibition trials, one study found greater left caudate nucleus activation in individuals who drank heavily versus individuals who drank light (Campanella et al., 2017), while during No-Go correct trials, individuals who drank heavily exhibited decreased activation in the putamen than individuals who drank light (Ahmadi et al., 2013). This discrepancy can likely be caused by the difference in contrast examined. Greater FC between the striatum (anterior putamen) and insula was found in individuals who use tobacco than in those who did not (Akkermans, 2019).

##### 3.2.2.5 Hippocampus

Two studies show lower hippocampal activation during inhibition. Lesser hippocampal activation during No-Go correct trials in individuals who drank heavily versus individuals who drank light (Ahmadi et al., 2013) and significantly lesser parahippocampal gyrus activation in both individuals who use tobacco and individuals who previously used tobacco versus controls during Stop trials (Nestor et al., 2011) was observed.

##### 3.2.2.6 Summary

During response inhibition, mixed findings were present in the ACC, striatum, and dlPFC, while consistently lesser activation was observed in the IFG and hippocampus. The IFG is especially important in inhibitory processes; thus, the finding of lesser rIFG activation provides a biomarker for unsuccessful inhibition in individuals using substances, though replication of this finding is needed.

#### 3.2.3 Negative Emotionality Domain

See **Table 3** for an overview of findings from this domain.

**Table 3.** Negative emotionality studies.

| Citation | Substance | Task Used | Experimental Group (n) | Age Mean (SD) | Percentage of Female participants (%) | Control Group (n) | Age Mean (SD) | Percentage of Female participants (%) | OFC | mPFC | ACC | Hippocampus | Amygdala |
| --- | --- | --- | --- | --- | --- | --- | --- | --- | --- | --- | --- | --- | --- |
| Brown-Rice et al., 2018 | Alcohol | Rank Emotion Task | 21 | 18-25 | 58% | 23 | 19-24 | 64% | - | ↑ | - | - | - |
| MacIlvane et al., 2020 | Alcohol | Emotional face-matching task | 39 | 39.90 (12.59) | 28% | 103 | 36.08 (11.14) | 55% | - | - | - | - | - |
| Gorka et al., 2020a | Alcohol | Threat-of-shock startle task | 38 | 23.8 (3.0) | 42% | 27 | 24.3 (2.8) | 44% | - | - | ↑ | - | - |
| Gorka et al., 2020b | Alcohol | Threat-of-shock startle task | 33 | 23.6 (2.8) | 39% | 24 | 24.0 (2.6) | 54% | - | ↓ | - | - | ↓ |
| Zhao et al., 2020 | Cannabis | Montreal imaging stress task | 28 | 25.54 (5.11) | 0% | 23 | 24.57 (3.55) | 0% | - | - | - | - | - |
| Onur et al., 2012 | Tobacco | Modified face perception task | 28 | 26.32 (2.79) | 50% | 28 | 26.88 (2.44) | 50% | - | - | - | - | - |

*Notes.* Abbreviations: SD = standard deviation; OFC = orbitofrontal cortex; mPFC = medial prefrontal cortex; ACC = anterior cingulate cortex

##### 3.2.3.1 mPFC

One study found greater middle frontal gyrus (MFG) activation in individuals who exhibited hazardous drinking patterns while presenting negative stimuli than individuals who did not exhibit hazardous drinking patterns (Brown-rice et al., 2018). Regarding FC findings, less mPFC-amygdala FC was associated with more binge episodes within the past 60 days and a younger age of AUD onset (Gorka et al., 2020).

##### 3.2.3.2 ACC

One study found greater dACC reactivity during uncertain and predictable threats in individuals diagnosed with AUD than those without AUD (Gorka et al., 2020).

##### 3.2.3.3 Amygdala

One study found no group differences between individuals with AUD and anxiety and individuals diagnosed with AUD only or healthy controls when examining fearful faces versus shapes or between individuals who use tobacco compared to individuals who do not smoke tobacco when examining emotional facial expressions (Onur et al., 2012).

##### 3.2.3.4 Summary

Greater middle frontal gyrus (Brown-Rice et al., 2018), lesser mPFC activation, and decreased amygdala-mPFC FC during unexpected threats were associated with more binge episodes in individuals who experience binge drinking episodes (Gorka et al., 2020). These findings follow studies suggesting lesser FC would suggest deficient control of amygdala activation by the mPFC. The finding of greater dACC activation in individuals diagnosed with AUD in response to threat than controls is also in line with previous data on the role of the dACC in contributing to increased anxious states in individuals with AUD (Gorka et al., 2020). Only one study found differences in the ACC in individuals diagnosed with AUD versus controls; more studies are needed to clarify the role of the ACC and other regions implicated in threat in individuals who use substances. This domain is especially important to explore as the ‘self-medication’ hypothesis has proposed extensive use of substances to alleviate anxiety or depression symptoms, leading to the ultimate development of a substance use problem.

## 4. Discussion

In this systematic review, we explored three neural pathways in individuals who use substances that define the neurofunctional domains suggested by Hien and colleagues (Hien et al., 2021). Our findings suggest altered activation and FC in reward, inhibition, and negative emotionality neural circuits in currently using, dependent on substances, or clinically diagnosed with SUD than controls. Notably, no studies on trauma-exposed individuals using, dependent on, or clinically diagnosed with SUD met the criteria for our systematic review. The implications of these findings are discussed below.

### 4.1 Reward Domain

Overall, the studies reviewed showed greater activation in individuals who use substances versus controls across many brain regions important in the mesocorticolimbic pathway, including the OFC, VTA, amygdala, and hippocampus, when using reward-related tasks such as cue-reactivity and monetary incentive tasks.

An increase in VTA and striatal activation during the presentation of cue-related activation is not surprising, given that the VTA is responsible for releasing dopamine to the striatum in response to rewarding stimuli (Cai & Tong, 2022). However, even in individuals with substance use dependence, there was still greater activation in these areas compared to individuals without substance use dependence to cue-related stimuli. This was surprising given previous literature that suggests individuals dependent on a substance no longer exhibit similar increases in dopaminergic initiation given the tolerance created for the substance (Koob & Volkow, 2016). We excluded intervention studies that gave participants a substance inside the scanner; thus, the contextual cues associated with that substance rather than the actual substance could provide a greater release of dopamine-producing a rewarding effect. We did find greater activation in the hippocampus in individuals who use substances versus controls, providing evidence of the importance of context cue processing during reward processing. Thus, these findings highlight that activation in response to viewing reward-related stimuli leads to greater activation of brain regions underlying reward even during dependence.

The increased activation in the VTA and striatal findings may suggest that when passively viewing rewards without the ability to receive the reward leads to an increase in reward-related activation, whereas actively receiving the reward or consciously making decisions with outcomes that will directly affect the participant may lead to deficient regulation of reward in the VTA, striatum, and ACC.

### 4.2 Inhibition Domain

Overall, studies found lesser activation in the IFG and hippocampus. In contrast, mixed findings in activation were present in the ACC, striatum, and dlPFC in individuals who use substances samples versus controls.

Given its important role in executing successful response inhibition, the findings of lesser IFG in individuals who use tobacco could correspond with the behavioral deficits seen in inhibiting their urge to use substances, even when the substance is causing functional difficulties in the person’s life. Deficits in the brain regions responsible for inhibition may lead to impulsive behavior associated with initiating and maintaining substance use (Kozak et al., 2019). More studies are needed to determine better the role of the ACC, striatum, and dlPFC in inhibitory processes.

### 4.3 Negative Emotionality Domain

The last domain is negative emotionality, whose phenotypical expression is related to the mesolimbic domain of the brain, encompassing the mPFC, ACC, and amygdala. Only 6 studies were eligible to be included in the review. Regardless, these results highlight greater dACC and lesser mPFC activation in individuals with hazardous drinking patterns and AUD compared to controls.

The dACC is important in the acquisition of conditioned fear. Thus, greater activation in the dACC could suggest that individuals that use substances have altered appraisal and expression of conditioned fear than controls. Interestingly, one study found greater MFG activation in response to negative images in individuals with hazardous drinking patterns versus individuals who do not exhibit hazardous drinking patterns (Brown-Rice et al., 2018). This finding is interesting, given the regulatory role of the MFG in responding to potentially threatening stimuli. However, the MFG is potentially trying to hyper-regulate regions responsible for fear expression (i.e., the amygdala), leading to maladaptive responses.

### 4.4 Trauma-exposed samples

No studies were identified that examined functional MRI correlates of reward salience, negative emotionality, or inhibition in substance use samples with trauma exposure that met the criteria of our PRISMA-guided systematic review. Given that trauma is a risk factor for the development of SU and SUD, as well as the large co-occurrence of PTSD and SUD it is likely that our sample of individuals who use substances has some proportion who were un-assessed trauma-exposed individuals. Therefore, studies are needed to examine the intersection of trauma history with functional correlates of reward salience, inhibition (or executive function), and negative emotionality in individuals who use substances. Determining how an individual responds to a stressor and how trauma-related factors may interact with neurobiological mechanisms to influence SU may facilitate screening individuals at risk and developing targeted clinical interventions to mitigate the escalation of substance use.

### 4.5 Limitations and Future Directions

There are limitations of the literature reviewed. First, this review highlights a lack of studies that have systematically measured trauma exposure in individuals who use substances when examining reward, inhibition, and negative emotionality neurocircuitry. We could not identify studies conducted in trauma-exposed samples; however, as noted above, it is plausible that participants in the studies where trauma exposure was not measured did indeed experience trauma, but trauma exposure was not assessed in these studies. It will be imperative to measure trauma exposure moving forward in studies using samples who use substances to start understanding the potential neural mechanisms associated with trauma that contribute to the development of substance use or vice versa. Second, while many studies examined neural correlates of reward-related neurocircuitry in SUDs, neuroimaging studies have yet to thoroughly explore the two other domains, inhibition, and negative emotionality, in individuals who use substances. Future studies should utilize tasks focusing on these domains to determine whether alterations are associated with substance use maintenance and development. Third of the studies reviewed, participants with varying levels of substance use were combined, including individuals who occasionally use, those who use heavily, and those diagnosed with SUD. More studies should focus on each level of the substance use cycle to understand the neural correlates of early use, transitioning to heavy use, and later disorder. Lastly, many studies reviewed contained small sample sizes (*n*’s<100 per group). In the future, studies utilizing bigger samples should be published to increase the power to determine effects.

There are also limitations related to our systematic review. First, our systematic review focuses on alcohol, tobacco, and cannabis use, given that these substances have been reported most in trauma studies (Bhalla et al., 2017). Other substances are important to explore, such as cocaine, heroin, and fentanyl. In addition, less research has been done on developing substance use post-trauma. Because our literature searches showed no studies examining trauma, we categorized the studies as not having measured trauma exposure. However, given that trauma exposure is a common risk factor for developing a SUD, such a sample is unlikely to be entirely unexposed to trauma. Therefore, the conclusions related to this sample should be interpreted with caution. More neuroimaging studies should be conducted to investigate the neural underpinnings associated with susceptibility to developing substance use problems post-trauma. Lastly, causality was not addressed in this review. It will be imperative to discover whether these findings result from using these substances or make an individual more prone to developing substance use issues. To answer this important question, longitudinal and twin samples should be used in the future.

### 4.6 Conclusion

This review systematically synthesizes neuroimaging studies focusing on three domains associated with substance use and may contribute to the overlapping mechanisms of SUD and PTSD – reward, inhibition, and negative emotionality. While more definitive research on individuals using substances, especially those with trauma exposure, and how these responses may contribute to the development and maintenance of substance use is needed, consistent with existing literature, we have highlighted deficits in each domain, specifically greater activation in regions involved in reward processing, lesser activation in regions involved in inhibition processing, and greater and lesser activation in regions involved in negative emotionality processes. Additionally, we propose that future studies should emphasize executive function and negative emotionality processes, investigate the role of trauma-related factors, and use longitudinal designs to better understand the underlying development of neurobiological alterations, providing mechanistic targets for preventative measures and treatment outcomes.

## References

Ahmadi, A., Pearlson, G. D., Meda, S. A., Dager, A., Potenza, M. N., Rosen, R., Austad, C. S., Raskin, S. A., Fallahi, C. R., Tennen, H., Wood, R. M., & Stevens, M. C. (2013). Influence of alcohol use on neural response to Go/No-Go task in college drinkers. Neuropsychopharmacology: Official Publication of the American College of Neuropsychopharmacology, 38(11), 2197–2208. https://doi.org/10.1038/npp.2013.119

Alderson Myers, A. B., Arienzo, D., Molnar, S. M., & Marinkovic, K. (2021). Local and network-level dysregulation of error processing is associated with binge drinking. NeuroImage. Clinical, 32, 102879. https://doi.org/10.1016/j.nicl.2021.102879

American Psychiatric Association. (2013). Diagnostic and statistical manual of mental disorders (5th ed.). https://doi.org/10.1176/appi.books.9780890425596

Ames, S. L., Wong, S. W., Bechara, A., Cappelli, C., Dust, M., Grenard, J. L., & Stacy, A. W. (2014). Neural correlates of a Go/NoGo task with alcohol stimuli in light and heavy young drinkers. Behavioural Brain Research, 274, 382–389. https://doi.org/10.1016/j.bbr.2014.08.039

Arias, A. J., Ma, L., Bjork, J. M., Hammond, C. J., Zhou, Y., Snyder, A., & Moeller, F. G. (2021). Altered effective connectivity of the reward network during an incentive-processing task in adults with alcohol use disorder. *Alcoholism*, Clinical and Experimental Research, 45(8), 1563–1577. https://doi.org/10.1111/acer.14650

Aron, A. R. (2007). The Neural Basis of Inhibition in Cognitive Control. The Neuroscientist, 13(3), 214–228. https://doi.org/10.1177/1073858407299288

Aupperle, R. L., Melrose, A. J., Stein, M. B., & Paulus, M. P. (2012). Executive Function and PTSD: Disengaging from Trauma. Neuropharmacology, 62(2), 686–694. https://doi.org/10.1016/j.neuropharm.2011.02.008

Baptiste, D.-L., Commodore-Mensah, Y., Alexander, K. A., Jacques, K., Wilson, P. R., Akomah, J., Sharps, P., & Cooper, L. A. (2020). COVID-19: Shedding light on racial and health inequities in the USA. Journal of Clinical Nursing, 29(15–16), 2734–2736. https://doi.org/10.1111/jocn.15351

Baxter, M. G., & Murray, E. A. (2002). The amygdala and reward. Nature Reviews Neuroscience, 3(7), Article 7. https://doi.org/10.1038/nrn875

Becker, A., Gerchen, M. F., Kirsch, M., Ubl, B., Subramaniapillai, S., Diener, C., Kuehner, C., Kiefer, F., & Kirsch, P. (2017). Frontostriatal connectivity during reward anticipation: A neurobiological mechanism cutting across alcohol use disorder and depression? Zeitschrift Für Psychologie, 225, 232–243. https://doi.org/10.1027/2151-2604/a000307

Becker, A., Kirsch, M., Gerchen, M. F., Kiefer, F., & Kirsch, P. (2017). Striatal activation and frontostriatal connectivity during non-drug reward anticipation in alcohol dependence. Addiction Biology, 22(3), 833–843. https://doi.org/10.1111/adb.12352

Berenz, E. C., Roberson-Nay, R., Latendresse, S., Mezuk, B., Gardner, C. O., Amstadter, A. B., & York, T. P. (2017). Posttraumatic Stress Disorder and Alcohol Dependence: Epidemiology and Order of Onset. *Psychological Trauma*_J*: Theory, Research*, Practice and Policy, 9(4), 485. https://doi.org/10.1037/tra0000185

Bergman, H. E., Przeworski, A., & Feeny, N. C. (2017). Rates of Subthreshold PTSD Among U.S. Military Veterans and Service Members: A Literature Review. *Military Psychology: The Official Journal of the Division of Military Psychology*, American Psychological Association, 29(2), 117–127. https://doi.org/10.1037/mil0000154

Bhalla, I. P., Stefanovics, E. A., & Rosenheck, R. A. (2017). Clinical epidemiology of single versus multiple substance use disorders. Medical Care, 55(9), S24–S32.

Bickel, W. K., Jarmolowicz, D. P., Mueller, E. T., Koffarnus, M. N., & Gatchalian, K. M. (2012). Excessive Discounting of Delayed Reinforcers as a Trans-Disease Process Contributing to Addiction and Other Disease-Related Vulnerabilities: Emerging Evidence. Pharmacology & Therapeutics, 134(3), 287–297. https://doi.org/10.1016/j.pharmthera.2012.02.004

Brown-Rice, K. A., Scholl, J. L., Fercho, K. A., Pearson, K., Kallsen, N. A., Davies, G. E., Ehli, E. A., Olson, S., Schweinle, A., Baugh, L. A., & Forster, G. L. (2018). Neural and psychological characteristics of college students with alcoholic parents differ depending on current alcohol use. Progress in Neuro-Psychopharmacology and Biological Psychiatry, 81, 284–296. https://doi.org/10.1016/j.pnpbp.2017.09.010

Bühler, M., Vollstädt-Klein, S., Kobiella, A., Budde, H., Reed, L. J., Braus, D. F., Büchel, C., & Smolka, M. N. (2010). Nicotine dependence is characterized by disordered reward processing in a network driving motivation. Biological Psychiatry, 67(8), 745–752. https://doi.org/10.1016/j.biopsych.2009.10.029

Burnette, E. M., Grodin, E. N., Ghahremani, D. G., Galván, A., Kohno, M., Ray, L. A., & London, E. D. (2021). Diminished cortical response to risk and loss during risky decision making in alcohol use disorder. Drug and Alcohol Dependence, 218, 108391. https://doi.org/10.1016/j.drugalcdep.2020.108391

Cai, J., & Tong, Q. (2022). Anatomy and Function of Ventral Tegmental Area Glutamate Neurons. Frontiers in Neural Circuits, 16, 867053. https://doi.org/10.3389/fncir.2022.867053

Chacon, N. C., Walia, N., Allen, A., Sciancalepore, A., Tiong, J., Quick, R., Mada, S., Diaz, M. A., & Rodriguez, I. (2021). Substance use during COVID-19 pandemic: Impact on the underserved communities. Discoveries, 9(4), e141. https://doi.org/10.15190/d.2021.20

Chen, H., Nebe, S., Mojtahedzadeh, N., Kuitunen-Paul, S., Garbusow, M., Schad, D. J., Rapp, M. A., Huys, Q. J. M., Heinz, A., & Smolka, M. N. (2021). Susceptibility to interference between Pavlovian and instrumental control is associated with early hazardous alcohol use. Addiction Biology, 26(4), e12983. https://doi.org/10.1111/adb.12983

Colombo, B., Bartesaghi, N., Simonelli, L., & Antonietti, A. (2015). The combined effects of neurostimulation and priming on creative thinking. A preliminary tDCS study on dorsolateral prefrontal cortex. Frontiers in Human Neuroscience, 9. https://www.frontiersin.org/articles/10.3389/fnhum.2015.00403

Cousijn, J., Wiers, R. W., Ridderinkhof, K. R., van den Brink, W., Veltman, D. J., Porrino, L. J., & Goudriaan, A. E. (2013). Individual differences in decision making and reward processing predict changes in cannabis use: A prospective functional magnetic resonance imaging study. Addiction Biology, 18(6), 1013–1023. https://doi.org/10.1111/j.1369-1600.2012.00498.x

Crane, N. A., Gorka, S. M., Weafer, J., Langenecker, S. A., de Wit, H., & Phan, K. L. (2017). Preliminary Evidence for Disrupted Nucleus Accumbens Reactivity and Connectivity to Reward in Binge Drinkers. *Alcohol and Alcoholism (Oxford*, Oxfordshire), 52(6), 647– 654. https://doi.org/10.1093/alcalc/agx062

Criaud, M., & Boulinguez, P. (2013). Have we been asking the right questions when assessing response inhibition in go/no-go tasks with fMRI? A meta-analysis and critical review. Neuroscience & Biobehavioral Reviews, 37(1), 11–23. https://doi.org/10.1016/j.neubiorev.2012.11.003

Dager, A. D., Anderson, B. M., Rosen, R., Khadka, S., Sawyer, B., Jiantonio-Kelly, R. E., Austad, C. S., Raskin, S. A., Tennen, H., Wood, R. M., Fallahi, C. R., & Pearlson, G. D. (2014). Functional magnetic resonance imaging (fMRI) response to alcohol pictures predicts subsequent transition to heavy drinking in college students. *Addiction (Abingdon*, England), 109(4), 585–595. https://doi.org/10.1111/add.12437

Daley, D. C. (2013). Family and social aspects of substance use disorders and treatment. Journal of Food and Drug Analysis, 21(4), S73–S76. https://doi.org/10.1016/j.jfda.2013.09.038

de Ruiter, M. B., Oosterlaan, J., Veltman, D. J., van den Brink, W., & Goudriaan, A. E. (2012). Similar hyporesponsiveness of the dorsomedial prefrontal cortex in problem gamblers and heavy smokers during an inhibitory control task. Drug and Alcohol Dependence, 121(1–2), 81–89. https://doi.org/10.1016/j.drugalcdep.2011.08.010

de Ruiter, M. B., Veltman, D. J., Goudriaan, A. E., Oosterlaan, J., Sjoerds, Z., & van den Brink, W. (2009). Response perseveration and ventral prefrontal sensitivity to reward and punishment in male problem gamblers and smokers. Neuropsychopharmacology_J: Official Publication of the American College of Neuropsychopharmacology, 34(4), 1027– 1038. https://doi.org/10.1038/npp.2008.175

Degenhardt, L., Bharat, C., Glantz, M. D., Bromet, E. J., Alonso, J., Bruffaerts, R., Bunting, B., de Girolamo, G., de Jonge, P., Florescu, S., Gureje, O., Haro, J. M., Harris, M. G., Hinkov, H., Karam, E. G., Karam, G., Kovess-Masfety, V., Lee, S., Makanjuola, V., … Wojtyniak, B. (2022). The associations between traumatic experiences and subsequent onset of a substance use disorder: Findings from the World Health Organization World Mental Health surveys. Drug and Alcohol Dependence, 240, 109574. https://doi.org/10.1016/j.drugalcdep.2022.109574

Duehlmeyer, L., Parsons, N., Malpas, C. B., & Hester, R. (2022). Functional connectivity during feedback learning in smokers. Addiction Biology, 27(1), e13109. https://doi.org/10.1111/adb.13109

Filbey, F. M., Dunlop, J., Ketcherside, A., Baine, J., Rhinehardt, T., Kuhn, B., DeWitt, S., & Alvi, T. (2016). FMRI study of neural sensitization to hedonic stimuli in long-term, daily cannabis users. Human Brain Mapping, 37(10), 3431–3443. https://doi.org/10.1002/hbm.23250

Gorka, S. M., Teppen, T., Radoman, M., Phan, K. L., & Pandey, S. C. (2020). Human Plasma BDNF Is Associated With Amygdala-Prefrontal Cortex Functional Connectivity and Problem Drinking Behaviors. International Journal of Neuropsychopharmacology, 23(1), 1–11. https://doi.org/10.1093/ijnp/pyz057

Grodin, E. N., Sussman, L., Sundby, K., Brennan, G. M., Diazgranados, N., Heilig, M., & Momenan, R. (2018). Neural Correlates of Compulsive Alcohol Seeking in Heavy Drinkers. Biological Psychiatry. Cognitive Neuroscience and Neuroimaging, 3(12), 1022–1031. https://doi.org/10.1016/j.bpsc.2018.06.009

Hien, D. A., López-Castro, T., Fitzpatrick, S., Ruglass, L. M., Fertuck, E. A., & Melara, R. (2021). A unifying translational framework to advance treatment research for comorbid PTSD and substance use disorders. Neuroscience & Biobehavioral Reviews, 127, 779– 794. https://doi.org/10.1016/j.neubiorev.2021.05.022

Hillman, K. L., & Bilkey, D. K. (2012). Neural encoding of competitive effort in the anterior cingulate cortex. Nature Neuroscience, 15(9), 1290–1297. https://doi.org/10.1038/nn.3187

Howard, J. D., & Kahnt, T. (2021). To be specific: The role of orbitofrontal cortex in signaling reward identity. Behavioral Neuroscience, 135(2), 210–217. https://doi.org/10.1037/bne0000455

Hu, S., Zhang, S., Chao, H. H., Krystal, J. H., & Li, C. R. (2016). Association of drinking problems and duration of alcohol use to inhibitory control in non-dependent young adult social drinkers. *Alcoholism*, Clinical and Experimental Research, 40(2), 319–328. https://doi.org/10.1111/acer.12964

Insel, T., Cuthbert, B., Garvey, M., Heinssen, R., Pine, D. S., Quinn, K., Sanislow, C., & Wang, P. (2010). Research domain criteria (RDoC): Toward a new classification framework for research on mental disorders. The American Journal of Psychiatry, 167(7), 748–751. https://doi.org/10.1176/appi.ajp.2010.09091379

Janes, A. C., Nickerson, L. D., Frederick, B. D. B., & Kaufman, M. J. (2012). Prefrontal and limbic resting state brain network functional connectivity differs between nicotine-dependent smokers and non-smoking controls. Drug and Alcohol Dependence, 125(3), 252–259. https://doi.org/10.1016/j.drugalcdep.2012.02.020

Kaysen, D., Dillworth, T. M., Simpson, T., Waldrop, A., Larimer, M. E., & Resick, P. A. (2007). Domestic violence and alcohol use: Trauma-related symptoms and motives for drinking. Addictive Behaviors, 32(6), 1272–1283. https://doi.org/10.1016/j.addbeh.2006.09.007

Khantzian, E. J. (1985). The self-medication hypothesis of addictive disorders: Focus on heroin and cocaine dependence. The American Journal of Psychiatry, 142(11), 1259–1264. https://doi.org/10.1176/ajp.142.11.1259

Khantzian, E. J. (1997). The self-medication hypothesis of substance use disorders: A reconsideration and recent applications. Harvard Review of Psychiatry, 4(5), 231–244. https://doi.org/10.3109/10673229709030550

Kilpatrick, D. G., Acierno, R., Resnick, H. S., Saunders, B. E., & Best, C. L. (1997). A 2-year longitudinal analysis of the relationships between violent assault and substance use in women. Journal of Consulting and Clinical Psychology, 65(5), 834–847. https://doi.org/10.1037//0022-006x.65.5.834

Kilpatrick, D. G., Acierno, R., Saunders, B., Resnick, H. S., Best, C. L., & Schnurr, P. P. (2000). Risk factors for adolescent substance abuse and dependence: Data from a national sample. Journal of Consulting and Clinical Psychology, 68(1), 19–30. https://doi.org/10.1037//0022-006x.68.1.19

Kilpatrick, D. G., Resnick, H. S., Milanak, M. E., Miller, M. W., Keyes, K. M., & Friedman, M. J. (2013). National estimates of exposure to traumatic events and PTSD prevalence using DSM-IV and DSM-5 criteria. Journal of Traumatic Stress, 26(5), 537–547. https://doi.org/10.1002/jts.21848

Kleinhans, N. M., Sweigert, J., Blake, M., Douglass, B., Doane, B., Reitz, F., & Larimer, M. (2020). FMRI activation to cannabis odor cues is altered in individuals at risk for a cannabis use disorder. Brain and Behavior, 10(10), e01764. https://doi.org/10.1002/brb3.1764

Koob, G. F., & Volkow, N. D. (2016). Neurobiology of addiction: A neurocircuitry analysis. The Lancet. Psychiatry, 3(8), 760–773. https://doi.org/10.1016/S2215-0366(16)00104-8

Kozak, K., Lucatch, A. M., Lowe, D. J. E., Balodis, I. M., MacKillop, J., & George, T. P. (2019). The neurobiology of impulsivity and substance use disorders: Implications for treatment. Annals of the New York Academy of Sciences, 1451(1), 71–91. https://doi.org/10.1111/nyas.13977

Kunas, S. L., Stuke, H., Heinz, A., Ströhle, A., & Bermpohl, F. (2022). Evidence for a hijacked brain reward system but no desensitized threat system in quitting-motivated smokers: An fMRI study. Addiction, 117(3), 701–712. https://doi.org/10.1111/add.15651

Kwako, L. E., Momenan, R., Litten, R. Z., Koob, G. F., & Goldman, D. (2016). Addictions Neuroclinical Assessment: A Neuroscience-Based Framework for Addictive Disorders. Biological Psychiatry, 80(3), 179–189. https://doi.org/10.1016/j.biopsych.2015.10.024

Lawn, W., Mithchener, L., Freeman, T. P., Benattayallah, A., Bisby, J. A., Wall, M. B., Dodds, C. M., Curran, H. V., & Morgan, C. J. A. (2020). Value-based decision-making of cigarette and nondrug rewards in dependent and occasional cigarette smokers: An FMRI study. Addiction Biology, 25(4), e12802. https://doi.org/10.1111/adb.12802

Lee, I., & Byeon, J. S. (2014). Learning-dependent Changes in the Neuronal Correlates of Response Inhibition in the Prefrontal Cortex and Hippocampus. Experimental Neurobiology, 23(2), 178–189. https://doi.org/10.5607/en.2014.23.2.178

Lessov-Schlaggar, C. N., Lepore, R. L., Kristjansson, S. D., Schlaggar, B. L., Barnes, K. A., Petersen, S. E., Madden, P. A. F., Heath, A. C., & Barch, D. M. (2013). Functional neuroimaging study in identical twin pairs discordant for regular cigarette smoking. Addiction Biology, 18(1), 98–108. https://doi.org/10.1111/j.1369-1600.2012.00435.x

Liddle, P. F., Kiehl, K. A., & Smith, A. M. (2001). Event-related fMRI study of response inhibition. Human Brain Mapping, 12(2), 100–109. https://doi.org/10.1002/1097-0193(200102)12:2<100::aid-hbm1007>3.0.co;2-6

Litten, R. Z., Ryan, M. L., Falk, D. E., Reilly, M., Fertig, J. B., & Koob, G. F. (2015). Heterogeneity of alcohol use disorder: Understanding mechanisms to advance personalized treatment. *Alcoholism*, Clinical and Experimental Research, 39(4), 579–584. https://doi.org/10.1111/acer.12669

MacIlvane, N., Fede, S. J., Pearson, E. E., Diazgranados, N., & Momenan, R. (2020). A Distinct Neurophenotype of Fearful Face Processing in Alcohol Use Disorder With and Without Comorbid Anxiety. *Alcoholism*, Clinical and Experimental Research, 44(11), 2212– 2224. https://doi.org/10.1111/acer.14465

Miyake, A., Friedman, N. P., Emerson, M. J., Witzki, A. H., Howerter, A., & Wager, T. D. (2000). The Unity and Diversity of Executive Functions and Their Contributions to Complex “Frontal Lobe” Tasks: A Latent Variable Analysis. Cognitive Psychology, 41(1), 49–100. https://doi.org/10.1006/cogp.1999.0734

Munakata, Y., Herd, S. A., Chatham, C. H., Depue, B. E., Banich, M. T., & O’Reilly, R. C. (2011). A unified framework for inhibitory control. Trends in Cognitive Sciences, 15(10), 453–459. https://doi.org/10.1016/j.tics.2011.07.011

Murphy, P., & Garavan, H. (2011). Cognitive predictors of problem drinking and AUDIT scores among college students. Drug and Alcohol Dependence, 115(1–2), 94–100. https://doi.org/10.1016/j.drugalcdep.2010.10.011

Nestor, L. J., McCabe, E., Jones, J., Clancy, L., & Garavan, H. (2018). Smokers and ex-smokers have shared differences in the neural substrates for potential monetary gains and losses. Addiction Biology, 23(1), 369–378. https://doi.org/10.1111/adb.12484

Nestor, L., McCabe, E., Jones, J., Clancy, L., & Garavan, H. (2011). Differences in “bottom-up” and “top-down” neural activity in current and former cigarette smokers: Evidence for neural substrates which may promote nicotine abstinence through increased cognitive control. NeuroImage, 56(4), 2258–2275. https://doi.org/10.1016/j.neuroimage.2011.03.054

Phelps, E. A. (2004). Human emotion and memory: Interactions of the amygdala and hippocampal complex. Current Opinion in Neurobiology, 14(2), 198–202. https://doi.org/10.1016/j.conb.2004.03.015

Pietrzak, R. H., Goldstein, R. B., Southwick, S. M., & Grant, B. F. (2011). Prevalence and Axis I comorbidity of full and partial posttraumatic stress disorder in the United States: Results from Wave 2 of the National Epidemiologic Survey on Alcohol and Related Conditions. Journal of Anxiety Disorders, 25(3), 456–465. https://doi.org/10.1016/j.janxdis.2010.11.010

Poorolajal, J., Haghtalab, T., Farhadi, M., & Darvishi, N. (2016). Substance use disorder and risk of suicidal ideation, suicide attempt and suicide death: A meta-analysis. *Journal of Public Health (Oxford*, England), 38(3), e282–e291. https://doi.org/10.1093/pubmed/fdv148

Schulte, M. T., & Hser, Y.-I. (2014). Substance Use and Associated Health Conditions throughout the Lifespan. Public Health Reviews, 35(2), https://web-beta.archive.org/web/20150206061220/http://www.publichealthreviews.eu/upload/pdf_files/14/00_Schulte_Hser.pdf. https://doi.org/10.1007/BF03391702

Schultz, W. (2015). Neuronal Reward and Decision Signals: From Theories to Data. Physiological Reviews, 95(3), 853–951. https://doi.org/10.1152/physrev.00023.2014

Seeley, W. W., Menon, V., Schatzberg, A. F., Keller, J., Glover, G. H., Kenna, H., Reiss, A. L., & Greicius, M. D. (2007). Dissociable Intrinsic Connectivity Networks for Salience Processing and Executive Control. Journal of Neuroscience, 27(9), 2349–2356. https://doi.org/10.1523/JNEUROSCI.5587-06.2007

Simmonds, D. J., Pekar, J. J., & Mostofsky, S. H. (2008). Meta-analysis of Go/No-go tasks demonstrating that fMRI activation associated with response inhibition is task-dependent. Neuropsychologia, 46(1), 224–232. https://doi.org/10.1016/j.neuropsychologia.2007.07.015

Sjoerds, Z., van den Brink, W., Beekman, A. T. F., Penninx, B. W. J. H., & Veltman, D. J. (2014). Cue reactivity is associated with duration and severity of alcohol dependence: An FMRI study. PloS One, 9(1), e84560. https://doi.org/10.1371/journal.pone.0084560

Strosche, A., Zhang, X., Kirsch, M., Hermann, D., Ende, G., Kiefer, F., & Vollstädt-Klein, S. (2021). Investigation of brain functional connectivity to assess cognitive control over cue-processing in Alcohol Use Disorder. Addiction Biology, 26(1), e12863. https://doi.org/10.1111/adb.12863

van Hell, H. H., Vink, M., Ossewaarde, L., Jager, G., Kahn, R. S., & Ramsey, N. F. (2010). Chronic effects of cannabis use on the human reward system: An fMRI study. European Neuropsychopharmacology: The Journal of the European College of Neuropsychopharmacology, 20(3), 153–163. https://doi.org/10.1016/j.euroneuro.2009.11.010

van Holst, R. J., Clark, L., Veltman, D. J., van den Brink, W., & Goudriaan, A. E. (2014). Enhanced striatal responses during expectancy coding in alcohol dependence. Drug and Alcohol Dependence, 142, 204–208. https://doi.org/10.1016/j.drugalcdep.2014.06.019

Volkow, N. D., & Blanco, C. (2021). Research on substance use disorders during the COVID-19 pandemic. Journal of Substance Abuse Treatment, 129, 108385. https://doi.org/10.1016/j.jsat.2021.108385

Wiers, C. E., Stelzel, C., Park, S. Q., Gawron, C. K., Ludwig, V. U., Gutwinski, S., Heinz, A., Lindenmeyer, J., Wiers, R. W., Walter, H., & Bermpohl, F. (2014). Neural correlates of alcohol-approach bias in alcohol addiction: The spirit is willing but the flesh is weak for spirits. Neuropsychopharmacology: Official Publication of the American College of Neuropsychopharmacology, 39(3), 688–697. https://doi.org/10.1038/npp.2013.252

Windle, M. (1994). Substance use, risky behaviors, and victimization among a US national adolescent sample. Addiction, 89(2), 175–182. https://doi.org/10.1111/j.1360-0443.1994.tb00876.x

Yalachkov, Y., Kaiser, J., & Naumer, M. J. (2009). Brain regions related to tool use and action knowledge reflect nicotine dependence. The Journal of Neuroscience_J: The Official Journal of the Society for Neuroscience, 29(15), 4922–4929. https://doi.org/10.1523/JNEUROSCI.4891-08.2009

Zhang, X., Chen, X., Yu, Y., Sun, D., Ma, N., He, S., Hu, X., & Zhang, D. (2009). Masked smoking-related images modulate brain activity in smokers. Human Brain Mapping, 30(3), 896–907. https://doi.org/10.1002/hbm.20552

Zhou, X., Zimmermann, K., Xin, F., Zhao, W., Derckx, R. T., Sassmannshausen, A., Scheele, D., Hurlemann, R., Weber, B., Kendrick, K. M., & Becker, B. (2019). Cue Reactivity in the Ventral Striatum Characterizes Heavy Cannabis Use, Whereas Reactivity in the Dorsal Striatum Mediates Dependent Use. Biological Psychiatry. Cognitive Neuroscience and Neuroimaging, 4(8), 751–762. https://doi.org/10.1016/j.bpsc.2019.04.006

